# ER-located PIN5 transporter generates potent auxin sinks driven by the IAA decarboxylation pathway

**DOI:** 10.1101/2024.01.24.576992

**Authors:** Milada Čovanová, Karel Müller, Nikoleta Klierová, Nayyer Abdollahi Sisi, Petr Skůpa, Ondřej Smetana, Petre Ivanov Dobrev, Kamil Růžička, Jan Petrášek

## Abstract

Auxin is an essential and well-investigated regulator of plant development. Still, many aspects determining its (sub-)cellular distribution, as well as its metabolic turnover, are obscure. PIN5 is a transporter that resides on the endoplasmic reticulum and is presumed to influence internal auxin homeostasis by direct sequestration and subsequent degradation. Due to its distinct expression pattern and incomplete metabolomics analyses, the exact role of PIN5 protein and the identity of downstream auxin degradation products show significant gaps. To this end, we utilized morphologically homogeneous tobacco BY-2 cell cultures. We show that the expression of *Arabidopsis thaliana AtPIN5* in the BY-2 system phenocopies so-called auxin starvation defects. Moreover, we reveal that the activity of *At*PIN5 leads, in extreme cases, to the broad range of processes accompanying programmed cell death (PCD). Notably, based on the recently updated knowledge on auxin metabolism, we also show that a significant part of auxin metabolites downstream of the *At*PIN5 activity are part of the re-emerged auxin decarboxylation pathway. Taking together, we report the direct induction of PCD by auxin stimulus and propose the physiological framework of the auxin decarboxylation route.

## INTRODUCTION

The phytohormone auxin, or indole-3-acetic acid (IAA), is vital for plant growth and development. The pattern formation at the level of individual tissues and organs is achieved by the formation of local auxin maxima or minima, orchestrated by the cell-to-cell transport of IAA (Weijers *et al*., 2021). Among the mediators of auxin flow emerge the transporters of the PIN-formed (PIN) family. Showing a remarkable asymmetrical distribution on the plasma membrane (PM), they coordinate auxin fluxes across the tissues, thereby controlling the downstream developmental programs (Adamowski & Friml, 2015).

Besides the well-investigated auxin fluxes between cells, mediated by the PM-located PINs, another group of PIN proteins was described as residing at the endoplasmic reticulum (ER). The so-called ER-PINs regulate internal auxin levels by conveying auxin into the ER lumen for subsequent metabolization (Mravec *et al*., 2009; Ding *et al*., 2012). The metabolic products downstream of the ER-located PIN activity have been examined several times and with various degrees of consistency (Mravec *et al*., 2009; Ding *et al*., 2012; Abdollahi Sisi & Růžička, 2020). In addition, the pathways accompanying the IAA breakdown have recently undergone revision (Müller *et al*., 2021; Hayashi *et al*., 2021; Dobrev *et al*., 2023), highlighting the IAA decarboxylation as another relevant auxin degradation route (Bandurski *et al*., 1995; Dobrev *et al*., 2023). Thus, many aspects of the IAA homeostasis controlled by ER-PINs remain unclear (Abdollahi Sisi & Růžička, 2020; Ung *et al*., 2023a).

The connection between auxin and programmed cell death (PCD) has been repeatedly proposed (Kacprzyk *et al*., 2022). It was shown that auxin pulses produced from the cells undergoing PCD in the outermost layers of the root cap are required for priming lateral root primordia in the primary root meristem (Xuan *et al*., 2015, 2016). It was also revealed that auxin can suppress certain instances of PCD (Kerchev *et al*., 2013). It has even been hypothesized that auxin is a product of tryptophan degradation occurring in defunct cells (Sheldrake, 2021). Nevertheless, the authentic effect of auxin activity to trigger PCD has never been presented.

Cell-suspension cultures have proved to be an excellent model system, which enables studying the effects of growth substances directly and in a homogeneous manner (Petrášek & Zazímalová, 2006; Petrasek *et al*., 2006). Here, we explore the impact of conditional expression of the well-characterized ER-located *Arabidopsis thaliana At*PIN5 (Mravec *et al*., 2009) in the BY-2 tobacco suspension cultures. We comprehensively analyze the immediate IAA metabolites downstream of PIN5 activity and reveal the remarkably elevated presence of products of the IAA decarboxylation pathway (Dobrev *et al*., 2023). We also describe the corresponding phenotypic consequences, including the extreme instances of PCD.

## RESULTS

### Inducible expression of *At*PIN5 in BY-2 cells phenocopies the symptoms of auxin starvation

The growth and proliferation of tobacco BY-2 cells depends on the exogenous supply of auxin to the media. Its lack is characterized by enhanced cell elongation and lowered cell division, termed collectively as symptoms of auxin starvation (Winicur *et al*., 1998). We presented earlier that interfering with auxin transport leads to similar morphological defects. The induction of the transgene encoding a PM-located *Arabidopsis thaliana At*PIN7 auxin carrier enhances cell elongation and division arrest, consistent with that occurring on the auxin-free medium (Fig. 1, A to C and Supplemental Fig. S1C; Mravec *et al*., 2008; Müller *et al*., 2019). We observed similar auxin-deficient phenotypes by perturbing the nuclear auxin signaling by the inducible expression of the stabilized version of the auxin signaling repressor *At*IAA17, *At*axr3-1 (Leyser *et al*., 1996; Knox *et al*., 2003; Figs. 1D-F). We were also able to pharmacologically reduce cell proliferation by the dose-dependent application of IAA antagonist PEO-IAA, which competitively binds the TIR1/AFBs auxin receptors (Nishimura *et al*., 2009; Fig. S1B). This indicates that the auxin starvation phenotypes are dependent on the TIR1/AFB signaling, in addition to the availability of auxin in the cells.

**Figure 1.**
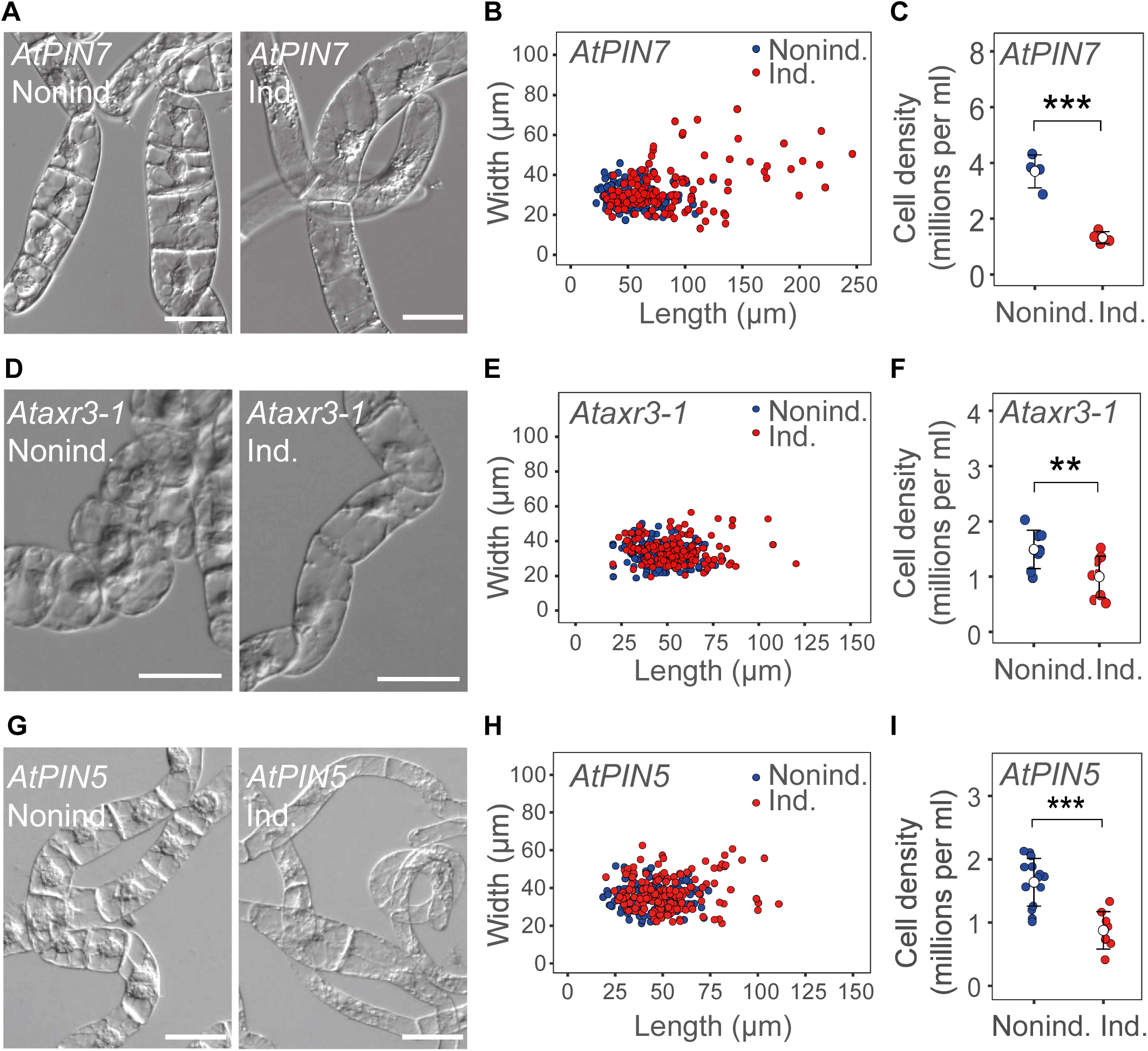
The conditional expression of *XVE::AtPIN5* in BY-2 cells leads to the auxin starvation phenotypes. **A** to **C)** The induction of the *XVE::AtPIN7* transgene results in increased cell elongation **A)**, plotted on **B)** (*P* < 0.001 by Student’s *t*-test), and decreased growth, monitored by the cell density in the media **C)**. **D** to **F)** The induction of *XVE::Ataxr3-1* phenocopies the symptoms of auxin starvation documented by representative image on **D)**, quantification of increased cell elongation (*P* < 0.001 by Student’s *t*-test) **E)** and reduced proliferation **F)**. **G** to **I)** The induction of *XVE::AtPIN5* in the cell line cultured in a standard medium with 2,4-D leads to increased cell elongation **G)**, plotted on **H)** (*P* < 0.05 by Student’s *t*-test), and reduced proliferation rate **I)**. On the dot plots, the open symbols are averages ± S.E. \*\**P* < 0.01, \*\*\**P* < 0.001 by Student’s *t*-test. Scale bars: 50 μm.

Next, to study the impact of internal auxin fluxes on the cell culture phenotypes, we expressed in the BY-2 cells the characterized *Arabidopsis thaliana* gene *AtPIN5* (Mravec *et al*., 2009; Abdollahi Sisi & Růžička, 2020). We employed two independent transcription inducible systems, XVE (Zuo *et al*., 2000; Petrasek *et al*., 2006) and GVG (Aoyama & Chua, 1997; Mravec *et al*., 2008), triggering the expression of the transgene at the beginning of the subculture interval. Upon the induction of *At*PIN5, the cells became remarkably expanded (Fig. 1G and 1H; Supplemental Fig. S1D and S1E), and their cell division reduced (Fig. 1I and Supplemental Fig. S1F), akin to the auxin starvation phenotypes (Fig. 1 A to F and Supplemental Fig. S1C). In addition, the auxin-deficient phenotypes caused by the ER-located *At*PIN5 corresponded to the lowered expression of the auxin reporter *DR5::GFP* (Supplemental Fig. S1G; see also Barbez *et al*., 2013). This indicates that the *At*PIN5 activity is capable of changing the internal availability of auxin to control the growth and expansion of the BY-2 cells.

### IAA enhances the symptoms of *At*PIN5-induced auxin starvation

The dependence of the BY-2 tissue cultures on the addition of the synthetic auxin (2,4-dichlorophenoxyacetic acid or 2,4-D) to the media is ascribed to the low levels of endogenous IAA in the cells, caused by its rapid metabolization (Simon *et al*., 2013; Seifertová *et al*., 2014; Müller *et al*., 2021). Accordingly, substituting 2,4-D with IAA also leads to auxin starvation phenotypes (Simon *et al*., 2013; Fig. 2A). To test the role of internal auxin fluxes under conditions where only native auxin is present, we cultivated the cells expressing *AtPIN5* under the supply of IAA, too. The control lines proliferated less intensively and resembled the auxin-starved ones. Surprisingly, the induction of *At*PIN5 led to even stronger auxin starvation phenotypes (Fig. 2, B and C). Thus, it seems that the PIN5-like transporters can enhance rapid IAA turnover in BY-2 cells.

**Figure 2.**
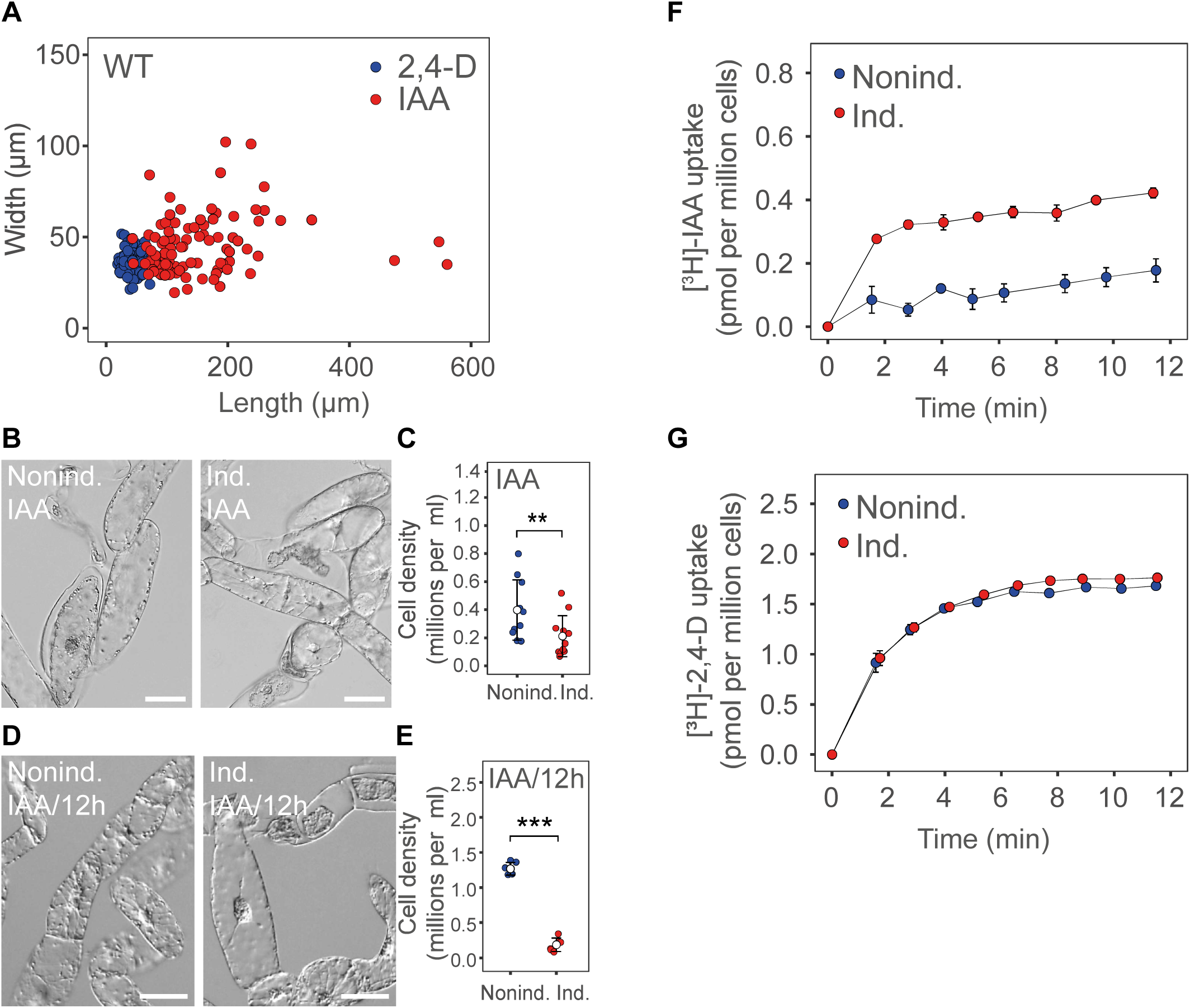
Replacing 2,4-D with its native analog IAA enhances the auxin starvation effects brought about by the induction of *XVE::AtPIN5*. **A)** The comparison of the cell proportions of the control *XVE::AtPIN5* cells cultivated in the medium supplemented with 2,4-D or IAA reveals the increase of the elongation growth in the presence of IAA (*P* < 0.001 by Student’s *t*-test). **B** to **C)** The proliferation rate of cells containing the induced *XVE::AtPIN5* transgene is even lower in the medium with IAA added at the beginning of subcultivation, the representative image on **B)**, the corresponding quantification of cell density on **C)**. **D** and **E)** The phenotype of the *XVE::AtPIN5* cells cultured in a medium continuously supplemented with IAA every 12 hours during the monitored cultivation period leads to the partial reversion of the starvation phenotypes in the control lines, however, the induction of the *XVE::AtPIN5* transgene leads to strong auxin starvation phenotypes, as documented at the representative image on **D)** and by the quantification of the cell growth on **E)**. **F** and **G)** The induction of *XVE::AtPIN5* results in higher uptake of [^3^H]-IAA **F)**, but shows no effect on uptake of [^3^H]-2,4-D **G)**. Scale bars: 50 μm. On the plots, the open symbols are averages ± S.E. \*\**P* < 0.01, \*\*\**P* < 0.001 by Student’s *t*-test.

To follow the original idea and to compensate for the fast IAA metabolization more efficiently, we were continuously adding IAA to the media every 12 h during the subcultivation interval. We observed no apparent signs of auxin deficiency (Fig. 2, D and E). Strikingly, though, the *AtPIN5* induction caused strong auxin starvation phenotypes (Fig. 2, D and E). This further underlines that PIN5 is able to generate a potent intracellular IAA sink.

ER-PINs are capable of transporting both IAA and 2,4-D (Ung *et al*., 2022). Hence, we measured the cellular accumulation of radioactively labeled [^3^H]-IAA and [^3^H]-2,4-D following the induction of *AtPIN5*. The lines showed a strong cellular uptake of [^3^H]-IAA (Fig. 2F). Interestingly, we observed no change in the *At*PIN5-mediated [^3^H]-2,4-D accumulation (Fig. 2G). Altogether, our data suggest that *At*PIN5 efficiently channels IAA from the cytosol to be delivered to likely highly active metabolic enzymes inside the ER lumen.

### Induction of *AtPIN5* promotes programmed cell death

To validate whether the strong auxin withdrawal from the cytosol by *At*PIN5 corresponds to a native downstream process, we analyzed the viability of the cells examined. The trypan blue exclusion test (van Wees, 2008), revealed an increased ratio of cells showing a dye-positive cytoplasm shrinkage as a sign of decreased cell viability following the *At*PIN5 induction (Fig. 3, A and B and Supplemental Fig. S2, A and B). Moreover, the intracellular trypan blue staining was even more apparent in the cells cultured on IAA (Fig. 3, A and B). Thus, the PIN5-mediated funneling of auxin from the cytosol can lead to increased mortality of BY-2 cells.

**Figure 3.**
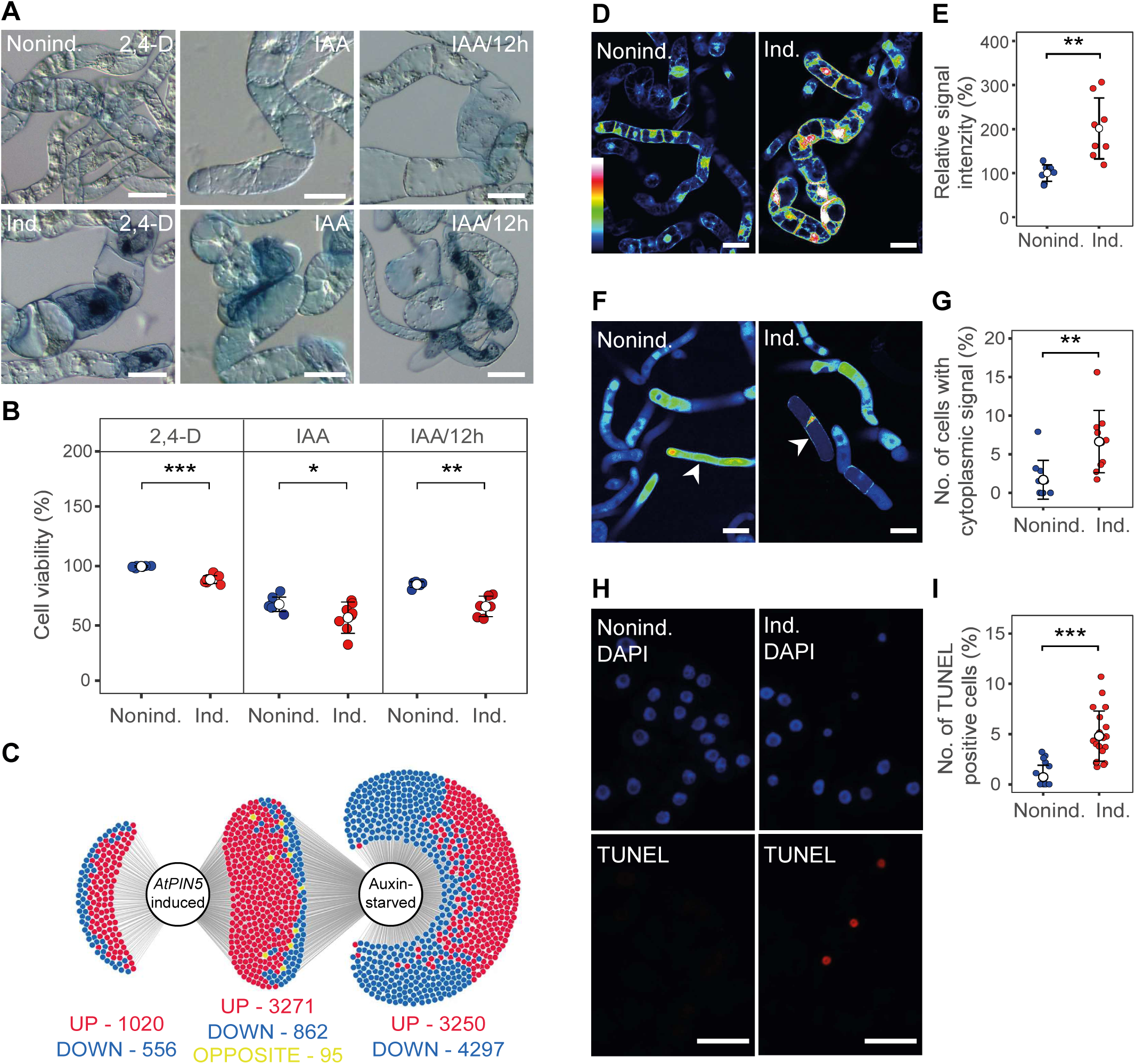
An increased incidence of symptoms of PCD following induction of *XVE::AtPIN5*. **A** and **B)** The replacement of 2,4-D with IAA leads to more frequent cytoplasm shrinkage and hereafter increased accumulation of the trypan blue dye inside cells (arrowheads), where *XVE::AtPIN5* was induced, as documented by representative images on **A)** and the inverse frequency of the trypan-blue intracellular staining on **B)**. **C)** DiVenn diagram indicating that a large part of differentially expressed genes following the *XVE::AtPIN5* induction is shared with those from the cultures undergoing auxin starvation; red, significantly upregulated genes; blue, downregulated genes; yellow, opposite expression changes. A single dot denotes a group of 10 genes. **D** and **E)** An increased fluorescence intensity on confocal images of the H2DCFDA staining of *XVE::AtPIN5* lines detects an elevated presence of ROS following the transgene induction **D)**, as quantified in **E)**. **F** and **G)** Confocal images of cells stained by BCECF-AM to mark acidic areas **F)**, as quantified in **G)**, document that the *XVE::AtPIN5* induction leads to decreased acidity in vacuoles but increased acidity in the cytoplasm and nucleus area (arrowheads). **H** and **I)** Representative images of TUNEL staining, marking the disintegration of nuclear DNA (red channel: TUNEL, blue channel: nuclear DAPI staining) **H)**, shows an enhanced labeling frequency in the lines where *XVE::AtPIN5* was induced **I)**. For **D)** and **F)**, the color intensity is indicated by the rainbow color lookup table (inset on **D)**). Scale bars: 50 μm. On the dot plots, the open symbols are averages ± S.E. \**P* < 0.05, \*\**P* < 0.01, \*\*\**P* < 0.001 by Student’s *t*-test.

Further, we analyzed the global transcriptional changes downstream of the *At*PIN5 activity to hint at the possible relevant physiological processes related to auxin starvation and lower cell viability. Indeed, using the PANTHER overrepresentation test (Mi *et al*., 2019), we found a remarkable overlap with the previously published transcriptional profiles from the BY-2 cells cultured in the auxin-free medium (Müller et al., 2021; Fig. 3C and Supplemental Fig. S2, C and D). This included pronounced changes in the expression of stress-related genes, including peroxidases (Supplemental Table S1). We further examined the transcriptional response exclusively linked with the *At*PIN5 expression (Supplemental Table S2), revealing a single enriched gene ontology term (GO:0000041) supported by 6 individual transcripts coding for transporters of transition metals (Peñarrubia *et al*., 2015). This suggests that the auxin starvation is associated with the enhanced expression of the stress-related peroxidases and that the *AtPIN5* expression leads to the enhanced metal ion transport, presumably activating the response preceding the cell death.

The increased expression of peroxidases following the *AtPIN5* induction suggests elevated activity of produced reactive oxygen species (ROS). The role of auxin has been implicated in the crosstalk with ROS homeostasis (Tognetti *et al*., 2012; Chen *et al*., 2014b). Hence, we utilized a sensor carboxy-2’,7’-dichloro-dihydro-fluorescein diacetate (H2DCFDA), which indicates the intracellular presence of various ROS, including H_2_O_2_ (Akter *et al*., 2021). Indeed, upon induction of *AtPIN5*, the cells displayed significantly increased H2DCFDA fluorescence, corroborating the enhanced ROS activity (Fig. 3, D and E and Supplemental Fig. S2E). PIN5 was previously shown to be a part of ER stress adaptive mechanisms (Chen *et al*., 2014a). Although we did not notice that the repeated additions of IAA into the media would influence the ER localization of *At*PIN5-GFP or lead to remarkable changes in the overall ER structural pattern (Supplemental Fig. S1G and S2F), it was earlier reported that pathways involving ROS production and ER stress are often preceding PCD (Petrov *et al*., 2015). This thereby further highlights that the lower viability of the *At*PIN5-GFP cells accompanied by ROS can perhaps be ascribed to the initiation of processes related to PCD.

Among the main features of PCD are vacuolar leakage and subsequent acidification of cytoplasm (Valandro *et al*., 2020). To this end, we utilized 2’,7’-Bis-(2-carboxyethyl)-5-(and-6)-carboxyfluorescein acetoxymethyl ester (BCECF/AM), a fluorescent dye used for analyzing lowered cytosolic pH as a mark of disrupted vacuolar integrity (Scheuring *et al*., 2015). Under control conditions, the cells contained several smaller vacuoles. The induction of *At*PIN5 led to the enlargement of central vacuoles, their occasional collapse and acidification of the whole cytoplasm and nucleus (Fig. 3, F and G). Finally, we tested the nuclear DNA fragmentation, a hallmark of PCD, using terminal deoxynucleotidyl transferase-mediated dUTP nick-end labeling (TUNEL) (Tripathi *et al*., 2017). The expression of *AtPIN5* (Fig. 3, H and I, and Supplemnetal Fig. S3A) resulted in an increased number of TUNEL-stained nuclei, indicating that DNA fragmentation takes part in the signaling cascade downstream of the rapid depletion of auxin in the cytoplasm. However, the decreased cell viability does not appear a general result of deficient auxin response. We also examined the TUNEL signal in cells expressing *AtPIN7* and *Ataxr3-1* in the medium supplemented with IAA in the 12-hour interval. We observed no relevant staining in any of these lines (Supplemental Fig. S3B).

We also tested whether the possibly disintegrated cell content can trigger (and enhance) the marks of auxin-induced starvation and PCD. We cultured wild-type BY-2 cells in a medium supplemented with the homogenized cell debris. The cell division rate remained unchanged (Supplemental Fig. S4A), and no obvious signs of increased cell mortality were observed (Supplemental Fig. S4B). This further evidences that the cell death induced by PIN5 expression is a result of the internal cellular activity, rather than being caused by any compound originating from disintegrated cells.

Taken together, we conclude that the observed decreased cell viability is associated with PCD, but not uncontrolled stress-related necrotic processes (Van Durme & Nowack, 2016), and it is a specific hallmark of PIN5-madiated auxin starvation.

### The activity of *At*PIN5 stimulates auxin degradation by the IAA decarboxylation pathway

To get insight into how the ER-mediated IAA conversion affects internal auxin homeostasis in light of recent findings (Müller *et al*., 2021; Hayashi *et al*., 2021; Dobrev *et al*., 2023; Fig. 4A), we have tracked the expanded list of IAA metabolites by liquid chromatography-mass spectrometry (LC-MS) following the *AtPIN5* induction. At the time point of adding IAA into media (*t*_0_), its degradation products were barely detectable and their amounts showed negligible differences between the mock and chemical inducer-treated cells (Supplemental Fig. S5A). 2 hours later, we observed that IAA was converted predominantly to amino-acid conjugates of IAA (Fig. 4B; see also Seifertová *et al*., 2014; Müller *et al*., 2021; Dobrev *et al*., 2023). The induction of *AtPIN5* usually led to a further increase in IAA conversion in this experimental setup (Fig. 4B and Supplemental Fig. S5B). These measurements illustrate rapid catabolism of IAA in BY-2 cells and evidence that the majority of detected metabolic compounds indeed originate from the supplied IAA.

**Figure 4.**
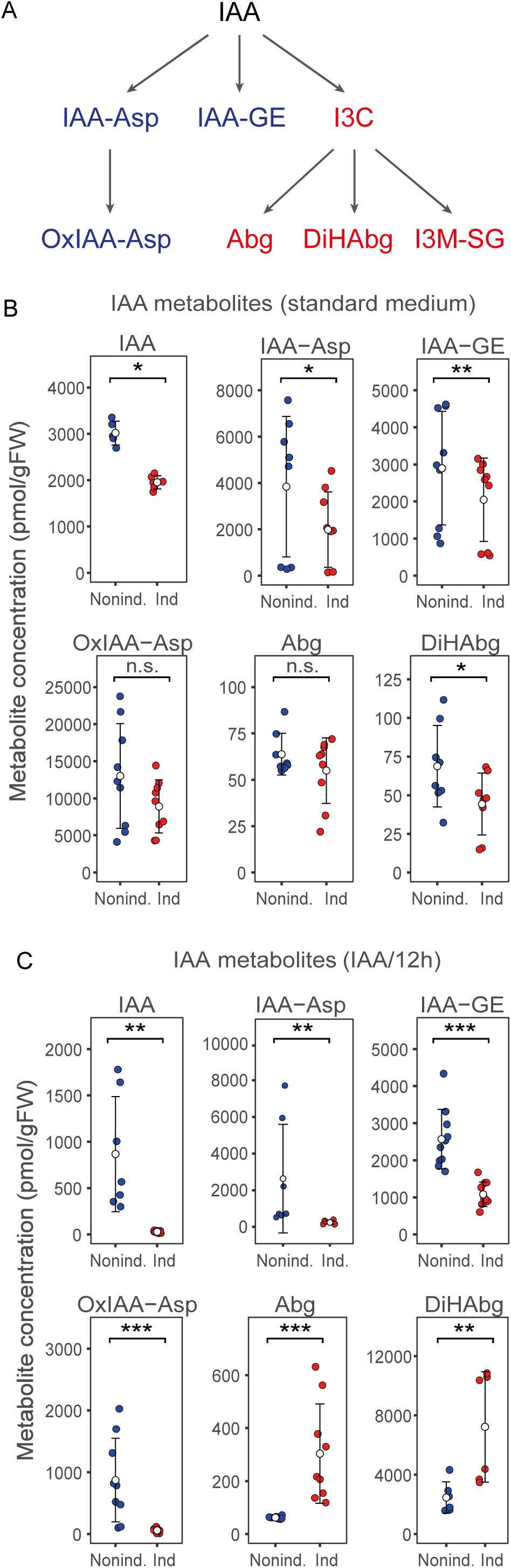
Inducible *XVE::AtPIN5* expression in BY-2 cells promotes IAA deactivation by the decarboxylation pathway. **A)** A scheme of current understanding of main IAA degradation pathways (products of IAA oxidation pathway in blue, IAA decarboxylation in red). **B)** LC-MS quantification of IAA and its metabolites in the cells expressing *XVE::AtPIN5* cultured on the standard 2,4-D-supplemented media shows a moderate decrease of IAA and its most degradation products. **C)** The induction of *XVE::AtPIN5* in the cells cultivated in media with IAA added every 12 hours leads to a dramatic drop of IAA and its conjugates, while the compounds belonging to the IAA decarboxylation pathway show elevated levels. On **B)** and **C)**, the open symbols indicate the mean from three independent measurements; the bars are ± S.E. \**P* < 0.05, \*\**P* < 0.01, \*\*\**P* < 0.001, by Student’s *t*-test, n. s.: not significant at *α*_0.05_. IAA: indole-3-acetic acid, IAA-Asp: IAA-aspartate, IAA-GE: IAA-glucose ester, OxIAA-Asp: Oxo-IAA-aspartate, Abg: ascorbigen, DiHAbg: dihydroascorbigen, I3M-SG: indole-3-methylglutathione. The cells were harvested 2 hours after the addition of 1 µM IAA.

Next, we quantified the IAA breakdown in the cells cultivated in a media supplied with IAA every 12 hours from the beginning of the subcultivation interval. 2 hours after addition, IAA was converted in the control lines into its conjugates as well (Fig. 4C). Surprisingly, the prior induction of *AtPIN5* dramatically reduced the internal IAA levels to close to trace amounts. Although we were also able to detect the activity of the common oxidative IAA degradation pathway, we found remarkably increased conversion towards IAA decarboxylation in these lines (Fig. 4C), as documented by the elevated levels of ascorbigen (Abg) and dihydroascorbigen (DiHAbg) (Dobrev *et al*., 2023; Fig. 4C). We therefore conclude that the PIN5-mediated auxin sinks are at least partially accompanied by the increased activity of the auxin decarboxylation pathway.

### The exogenous application of I3C raises the symptoms of cell death in BY-2 cells

It has been proposed earlier that indole-3-carbinol (I3C), an unstable immediate IAA decarboxylation product and precursor of Abg (Fig. 4A) (Hrnčiřík *et al*., 1998; Chhajed *et al*., 2020; Dobrev *et al*., 2023), can function as an auxin antagonist and can stimulate vacuolar disintegration and processes related to autophagy in *Arabidopsis* in consequence of the defense reaction (Katz *et al*., 2015; Katz & Chamovitz, 2017). We therefore incubated the cells with I3C to relate its role to the PIN5-mediated programmed cell death in the BY-2 cultures. Despite the default presence of 2,4-D in the media, I3C treatment affected the cell expansion and proliferation rates (Fig. 5, A and B). I3C also slightly lowered their overall viability (Fig. 5, C and D). However, it remarkably increased the frequency of PCD, as documented by TUNEL staining (Fig. 5, E and F). Taken together, we propose that ER-resided PINs are capable of generating powerful IAA sinks to activate IAA decarboxylation activity and, hereafter, pathways leading to PCD.

**Figure 5.**
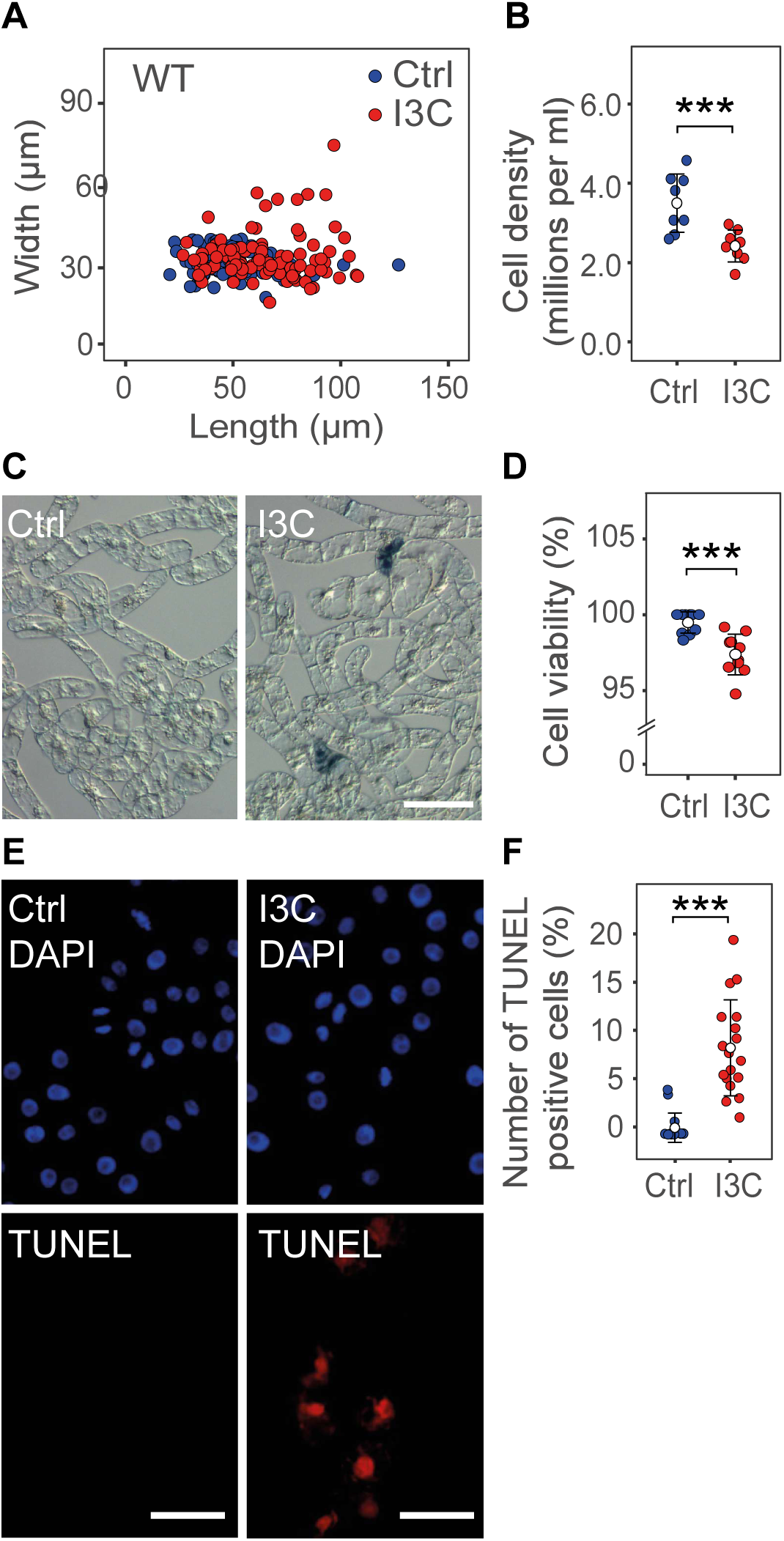
The I3C treatment leads to growth arrest and reduced viability of BY-2 cells. **A)** The addition of 50 µM I3C to the media leads to increased elongation of BY-2 cells (*P* < 0.001 by Student’s *t*-test) and **B)** decreased proliferation, monitored by the cell density in the media. **C** to **F)** The I3C treatment also results in increased cell mortality monitored by trypan blue **C** and **D)** and TUNEL staining (red channel: TUNEL, blue channel: nuclear DAPI staining) **E** and **F)**. Scale bars: 50 μm. On the dot plots, the open symbols are averages ± S.E. \*\*\**P* < 0.001 by Student’s *t*-test.

## DISCUSSION

### A possible mechanism underlying PIN5-mediated auxin sinks

The classical concept of polar auxin transport has premised the existence of efflux carriers asymmetrically localized at PM as the central prerequisite for the directional flow of IAA (Rubery & Sheldrake, 1974; Raven, 1975). Later, the PIN auxin transporters were discovered, matching henceforth the proposed attributes of the main IAA carriers (Gälweiler *et al*., 1998; Petrasek *et al*., 2006). ER-located PINs were found subsequently, and the relevant works highlighted their capability to influence internal auxin homeostasis (Mravec *et al*., 2009; Ding *et al*., 2012; Abdollahi Sisi & Růžička, 2020). Further, the detailed crystallographic studies evidenced that the transporting mechanism employed by both PM and ER-located PINs is identical (Su *et al*., 2022; Ung *et al*., 2022; Yang *et al*., 2022).

Yet, many aspects of the PIN-mediated transport remain obscure. Compelling arguments supported by the structural models suggest that IAA is transported from the cytosol against concentration gradients at the expense of membrane potential (Su *et al*., 2022; Ung *et al*., 2022, 2023a,b; Yang *et al*., 2022; Andersen *et al*., 2023). Indeed, in the root apical meristem, PM-located PINs concentrate auxin in the uttermost root tip (Adamowski & Friml, 2015). They also establish and maintain auxin maxima, determining the position of flower primordia in the shoot apical meristem in an equivalent manner (Reinhardt *et al*., 2003). In contrast, the canalization hypothesis proposes that auxin is funneled away from the site of its external application, i.e., source, towards the sink in the established vascular tissue determining the connective vascular strands (Mitchison, 1980; Sachs, 1991; Ravichandran *et al*., 2020). Similarly, the root or hypocotyl tropic reactions depend on the transport outwards from the tissue with higher auxin concentration (Friml *et al*., 2002; Ottenschläger *et al*., 2003; Rakusová *et al*., 2011; Band *et al*., 2012). The expression of PIN5 proteins leads to a remarkable auxin deficiency in BY-2 cells. In the conditions when 2,4-D does not interfere with the internal IAA fluxes (Fig. 5B), the excess of IAA leads to even enhanced PIN5-mediated auxin degradation rates. Hence, in this system, the PIN5-like auxin transporters likely operate in the source-sink mode, too, perhaps by the mechanism of facilitated diffusion, as earlier presumed for auxin canalization (Mitchison, 1980; Rolland-Lagan & Prusinkiewicz, 2005).

### Auxin metabolization as a prompt for PCD

Along with the auxin-dependent processes, PCD is an inherent part of plant development. The connections between auxin and PCD pathways have been proposed, but mostly indirectly (reviewed in Van Durme & Nowack, 2016; Sheldrake, 2021; Kacprzyk *et al*., 2022). A chemical screen performed by Kerchev *et al*. (2013) revealed that the H_2_O_2_-dependent cell death caused by defective photorespiration can be reverted by both synthetic auxins and IAA. Auxin was demonstrated as a central regulator of xylem patterning, where PCD gives rise to conductive vessels (Escamez & Tuominen, 2014; Ruzicka *et al*., 2015); direct transdifferentiation of the tissue cultures into xylem vessels can also be induced by external cues (Fukuda & Komamine, 1980; Kubo *et al*., 2005; Ruzicka *et al*., 2015). In addition, various stages of lateral root development are determined by auxin activity in parallel to PCD (Casimiro *et al*., 2001; Benková *et al*., 2003; Swarup *et al*., 2008; Escamez *et al*., 2020). Auxin serves as a signal produced by the cells undergoing PCD in the root cap to prime the prospective lateral roots in the primary root meristem (Xuan *et al*., 2015, 2016). Ultimately, Di Mambro et al. (2019) presented the model that the activity of *PIN5* in the lateral root cap affects root meristem size, but the direct influence of PIN5 on PCD in the lateral root cap has not been investigated.

We show that the ER-mediated auxin sinks can trigger PCD in BY-2 cultures. We have not noticed any cells resembling forming xylem vessels following the *AtPIN5* induction. Instead, we report phenotypes associated with common PCD and present lines of evidence that they are linked with the auxin decarboxylation activity. I3C is among native metabolites of glucosinolates, a diverse group of small compounds produced predominantly in *Brassicales*, functioning as agent in the insect defense (Fahey *et al*., 2001). It is also a product of auxin decarboxylation pathway in a multitude of the (non-)*Brassicales* plant species (Bandurski *et al*., 1995; Kerk *et al*., 2000; Dobrev *et al*., 2023). I3C can induce some symptoms of PCD in *Arabidopsis*, in a competitive manner to IAA (Katz *et al*., 2015; Katz & Chamovitz, 2017). Hence, our data suggest that the two pathways determining PCD overlap and share an unexpectedly common and likely evolutionally conserved functional output.

### IAA decarboxylation pathway downstream of PIN5-mediated auxin sequestration

PIN5 transports auxin into the ER in order to promote auxin degradation (Mravec *et al*., 2009; Ung *et al*., 2023a). The levels of IAA-aspartate and IAA-glutamate conjugates are elevated following *At*PIN5 overexpression (Mravec *et al*., 2009). Paradoxically, the GRETCHEN HAGEN (GH3) enzymes catalyzing the IAA conjugation are generally seen in the cytoplasm (Ludwig-Müller *et al*., 2009; Ostrowski *et al*., 2014; Di Mambro *et al*., 2019; Abdollahi Sisi & Růžička, 2020; Zou *et al*., 2022). We have recently uncovered a new level of complexity of the IAA metabolome. Among known IAA breakdown routes emerges the auxin decarboxylation pathway, which includes Abg and DiHAbg. Even if occurring across angiosperms, the purpose of auxin decarboxylation is unknown (Dobrev *et al*., 2023). Strikingly, we found that the relevant metabolic compounds are related to increased AtPIN5 activity. Hence, we conclude that IAA decarboxylation is the main and potent auxin degradation pathway downstream of PIN5.

## EXPERIMENTAL PROCEDURES

### Plant material, growth conditions and treatments

The tobacco cell lines BY-2 (*Nicotiana tabacum* L., cv. Bright Yellow-2; Nagata *et al*., 1992) were standardly cultivated in a liquid medium supplemented with 1 µM 2,4-D as described in Petrasek *et al*. (2006) unless stated otherwise. The transformation of cell lines, RT-qPCR validation of the transgene expression, and cell line maintenance were done as described earlier (Petrasek *et al*., 2006; Müller *et al*., 2021). By rule, the cells in the exponential growth stage were used, typically 4-d-old, unless specified otherwise.

The expression of the *AtPIN7, AtPIN5* and *Ataxr3-1* genes was induced by 1 µM dexamethasone (30 mM stock solution, GVG lines) or 3 µM β-estradiol (30 mM stock solution, XVE lines) at the beginning of the subculture interval. In addition, 2-(1H-indol-3-yl)-4-oxo-4-phenyl-butyric acid (PEO-IAA; OlChemIm, Olomouc, Czech Republic, 10 mM stock solution) and indole-3-carbinol (I3C; Sigma-Aldrich, Merck KGaA, Darmstadt, Germany, 160 mM stock solution) were used. The respective compounds were applied to the cells at the beginning of the subculture interval, and the corresponding amount of the solvent (DMSO) was added to the controls.

### Molecular transgenic work

The dexamethasone-inducible gene constructs, *35S>>GVG::AtPIN5* and *35S>>GVG::AtPIN7*, including the corresponding cell cultures, have been described earlier (Petrasek *et al*., 2006; Mravec *et al*., 2009). For creating *G10-90>>XVE::AtPIN5*, the corresponding *AtPIN5* open reading frame was cloned with the Gateway system (Invitrogen, Thermo Fisher Scientific, Waltham, MA, USA) into the pMDC7 vector (Curtis & Grossniklaus, 2003). The respective Gateway entry clones were generated either by the amplification of wild-type DNA template or that originating from the *PIN5-GFP^MGS^* plants (Verna *et al*., 2015) (primer sequences AAAAAGCAGGCTATATGATAAATTGTGGAGAT and AAGAAAGCTGGGTTTCAATGAATAAACTCC). The *XVE::AtPIN5-GFP*/*DR5::RFP* cell line was obtained by transformation of the *XVE::AtPIN5-GFP* line with the *DR5::RFP* construct (Barbez *et al*., 2013). The *G10-90>>XVE::Ataxr3-1* line was generated by the transformation of BY-2 cells with the construct kindly provided by Jiří Friml.

### Microscopy and image analysis

Nomarski differential interference contrast (DIC) microscopy was performed with the Nikon Eclipse E600 microscope (Nikon, Tokyo, Japan) equipped with a color digital camera (DVC 1310C, Austin, Texas, USA). Cell length and diameter were measured manually using the Fiji image analysis software (Rueden *et al*., 2017). 200 cells in each data set were measured. Cell density was determined by counting the cells with the Fuchs-Rosenthal hemocytometer slide, typically in the fourth day of subcultivation or periodically during the entire period; individual values represent the average of at least six aliquots of every sample. For the fluorescence imaging, Zeiss LSM900 laser scanning microscope (Axio Observer 7, inverted) with Airyscan 2 with Multiplex mode (Carl Zeiss, Jena, Germany) with appropriate filter sets to visualize reporter genes GFP (excitation 488 nm and emission 509 nm wavelengths) and/or mRFP (excitation 590 nm and emission 599 nm wavelengths) was used. Detailed images were obtained with the 40x or 63x water/oil immersion objectives.

### Assays on cell viability, ROS content and PCD

The trypan blue exclusion test was used to visualize the penetration of the non-viable cells by the dye (van Wees, 2008). Approximately 10 μl of the 0.4 % trypan blue solution (Sigma-Aldrich) was added to the 0.5 ml cell suspension and recorded subsequently. Cell viability was determined as the percentage of dye-negative cells relative to the total number of cells.

For the detection of reactive oxygen species (ROS), the cells were incubated with the 10 μM H2DCFDA (2′,7′-dichlorfluorescein-diacetate; Thermo Fisher Scientific, Waltham, Massachusetts, USA) solution (Akter *et al*., 2021) for 15 min and washed twice with the BY-2 medium. The relative ROS accumulation is presented as the mean pixel intensity of cells determined by the Zeiss Zen software. In total, about four hundred cells were counted from three independent experiments. The integrity of the vacuoles was determined after 10 min of staining with 10 μM BCECF-AM ((2′,7′-bis-2-carboxyethyl)-5-(and-6) carboxyfluorescein; Sigma-Aldrich; Scheuring *et al*., 2015). The number of cells with aberrant vacuolar integrity was assessed as a ratio of cells with the cytoplasmic signal to that of the cells showing a normal vacuolar signal. About five hundred cells were processed in three independent experiments. In both cases, the confocal images were acquired with the Plan-Apochromat 10x/0.45 DIC M27 objective.

TUNEL (terminal deoxynucleotidyl transferase dUTP nick end labeling) assay (Tripathi *et al*., 2017) was performed in 2-day-old BY-2 cells according to the manufacturer’s protocol (TMR red in situ cell death detection kit, Roche Diagnostics GmbH, Heidelberg, Germany) with modifications as described earlier (Smetana *et al*., 2012). The labeled cells were examined using a color digital camera (DVC 1310C, Austin, Texas, USA) at the Nikon Eclipse E600 microscope. All cells were analyzed from at least 10 view fields, covering approximately 100 cells each. The quantification is expressed as a percentage of cells carrying the TUNEL signal divided by the total cell number in the visual field. Positive controls, where the samples were additionally treated with DNAse I to detect the positive TUNEL reaction, and negative controls, where the TdT enzyme was omitted from the procedure, were imaged alongside with every experiment.

To treat wild-type BY-2 cells with cell debris, one-quarter volume of the total 2-day-old cell culture was collected. The cells were ground with a pestle in a mortar in liquid nitrogen. The homogenized cell content was then washed out by the original medium, and the flowthrough was then returned to the growing cell culture. The cells were subsequently cultivated for two more days before examining their physiological properties.

### Auxin uptake assays

Auxin uptake assays were performed in 3-day-old cells as described earlier (Delbarre *et al*., 1996; Petrásek *et al*., 2003). [^3^H]-2,4-D (10 Ci.mmol^-1^) or [^3^H]-IAA (25 Ci.mmol^-1^) (American Radiolabeled Chemicals, Inc., St Louis, MO, USA) were added to the equilibrated cell suspension at the beginning of the accumulation assay to the final concentration of 2 nM. The presented values are means of each triplicate from a representative experiment out of three biological repetitions.

### IAA metabolic profiling

The harvesting of the 2-d-old BY-2 cells was done 2 hours after the addition of 1 µM IAA into the media, unless stated otherwise. Auxin metabolites were extracted according to previously described procedures (Dobrev & Kamínek, 2002; Dobrev *et al*., 2023), and determined by the LC-MS quantitative analysis further detailed in Dobrev *et al*. (2023). The samples were processed in triplicates. The results are presented as a mean of at least three independent biological repetitions.

### Transcriptome analysis

The total RNA was isolated using RNeasy Plant Mini kit (Qiagen, Düsseldorf, Germany) from 50-100 mg of cells cultured in medium for 48 h with IAA added every 12 h, 2 h after the last addition of IAA. Isolated RNA was treated with DNA-Free kit (Thermo Fisher Scientific). RNA purity, concentration and integrity were evaluated on 0.8% agarose gels (v/w) and with the Bioanalyzer RNA Nano 6000 Assay Kit (Agilent Technologies, Santa Clara, CA, USA). RNA sequencing was performed using Illumina NovaSeq 6000 platform. The analysis resulted in at least 15 million 150 bps read pairs. The raw reads were quality-filtered using Rcorrector and Trim Galore scripts (Song & Florea, 2015). Transcript abundances (transcripts per million – TPM) were determined using the Salmon software package (Patro *et al*., 2017) with parameters --posBias, --seqBias, --gcBias, --numBootstraps 30. The index was built from the *Nicotiana tabacum* v1.0 cDNA dataset (Edwards *et al*., 2017), by including the *AtPIN5* (*AT5G16530*) gene sequence. Visualization, quality control of data analysis and determination of differentially expressed genes were determined using the sleuth package in R (Love *et al*., 2014; Pimentel *et al*., 2017) (version 0.29.0). Transcripts with the q-value <= 0.05 and log_2_ fold change >=1 (upregulated) or <= -1 (down-regulated) were considered to be significantly differentially expressed. Other criteria for selection of gene list of interest are specified in corresponding result section.

Gene ontology analysis (statistical overrepresentation test) was done using the Panther Classification System (Mi *et al*., 2019). Raw sequencing data are available through GEO Series accession number GSE260945 at the Gene Expression Omnibus (https://www.ncbi.nlm.nih.gov/geo/). The grouping of differentially expressed genes was visualized with the DiVenn software (Sun *et al*., 2019).

## ACKNOWLEDGEMENTS

We thank Jiří Friml for kindly providing the *Ataxr3-1* construct, Enrico Scarpella for the *PIN5-GFP^MGS^* lines, Zuzana Vondráková and Nikola Drážná for technical assistance, and Kateřina Malínská for help with image analysis. The work was supported by the Czech Science Foundation (project 19-23773S) to KR, JP and PID. Microscopy was done at the Imaging Facility of the Institute of Experimental Botany and was supported by the Ministry of Education, Youth and Sports of the Czech Republic (LM2023050 Czech-Bioimaging).

## AUTHOR CONTRIBUTIONS

JP and KR conceived the research. JP, KR and MČ designed the experiments. MČ performed most experiments. KM did RT-qPCR and transcriptome analysis. NK and NAS prepared DNA constructs. PS provided preliminary data essential for the initiation of the study. OS performed part of the PCD assays. PID did metabolic profiling. KR, JP and MČ wrote the manuscript. KR acquired funding. All authors have exercised their right to comment on the manuscript.

## SUPPLEMENTAL FIGURE LEGENDS

**Figure S1.**
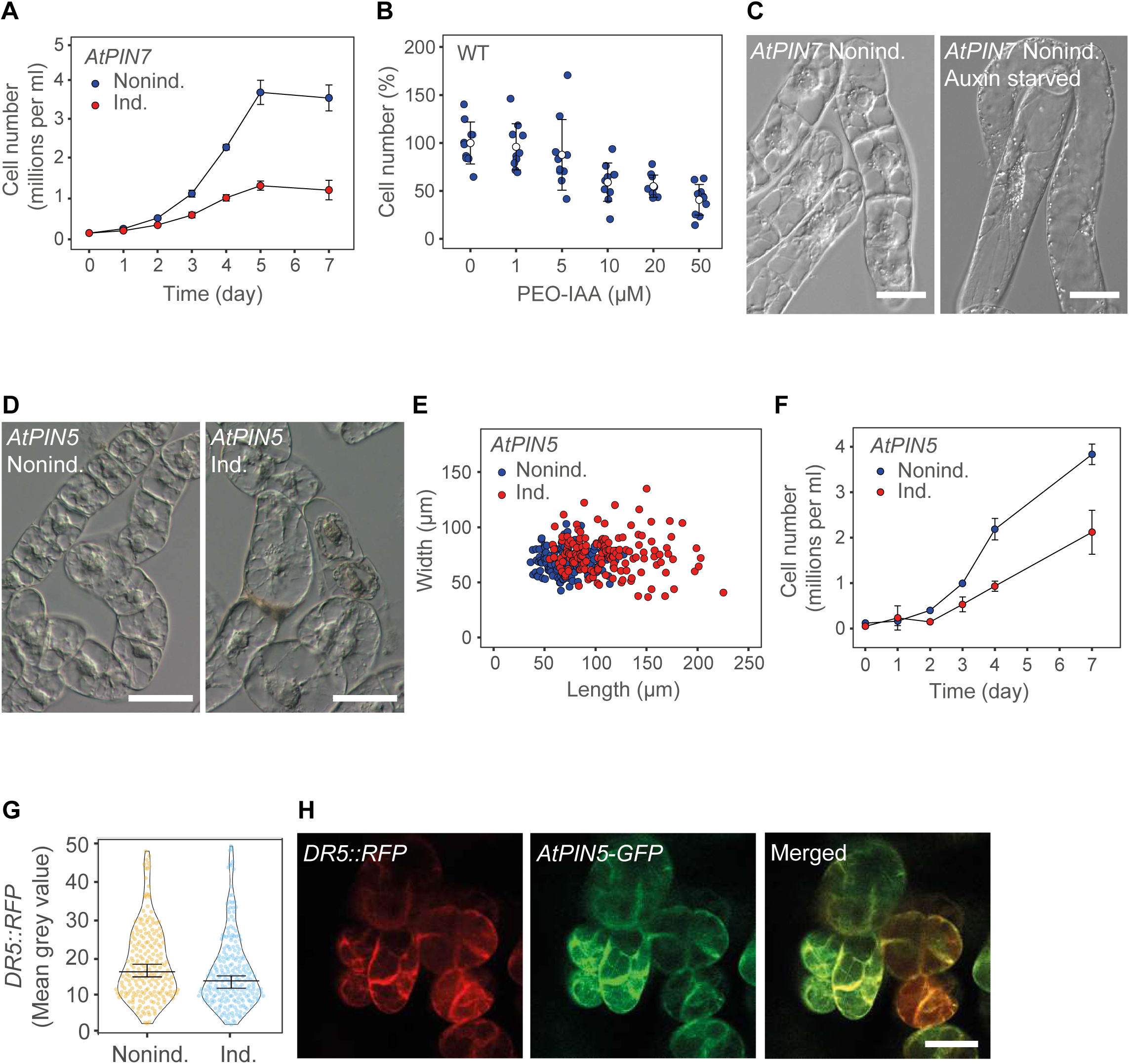
Additional aspects of auxin deficiency of tobacco BY-2 cells. **A)** The induction of the *GVG::AtPIN7* transgene leads to decreased cell density, reflecting their lower proliferation rate during the whole subcultivation interval. **B)** The cell density of the wild-type BY-2 cells decreases with the growing concentration of supplied PEO-IAA, an inhibitor of auxin-dependent gene expression (F(5,48) = [11.112], *P* < 0.001 by one-way ANOVA). **C)** The enhanced cell expansion is typical for cultivation on the auxin-free medium (shown on the *GVG::AtPIN7* line without the chemical inducer). **D** to **F)** The conditional expression of *GVG::AtPIN5* in the cell lines cultured in standard medium with 2,4-D leads to increased cell expansion **D** and **E)** (*P* < 0.001 by Student’s t-test) and reduced proliferation rates **F)**. **G** to **H)** The induction of *XVE::AtPIN5-GFP* in the cells co-expressing auxin *DR5::RFP* reporter results in a reduction of the fluorescence in cells in the upper-intensity quantiles **G)**, and PIN5-GFP colocalizes with the ER-targeted RFP **H)**. On **A)**, **B)** and **F)**, the open symbols are averages ± S.E. On **G)**, the vertical error bars represent median ± 95% confidence intervals. Scale bars: 50 μm on **C)**, **D)** and **H)**.

**Figure S2.**
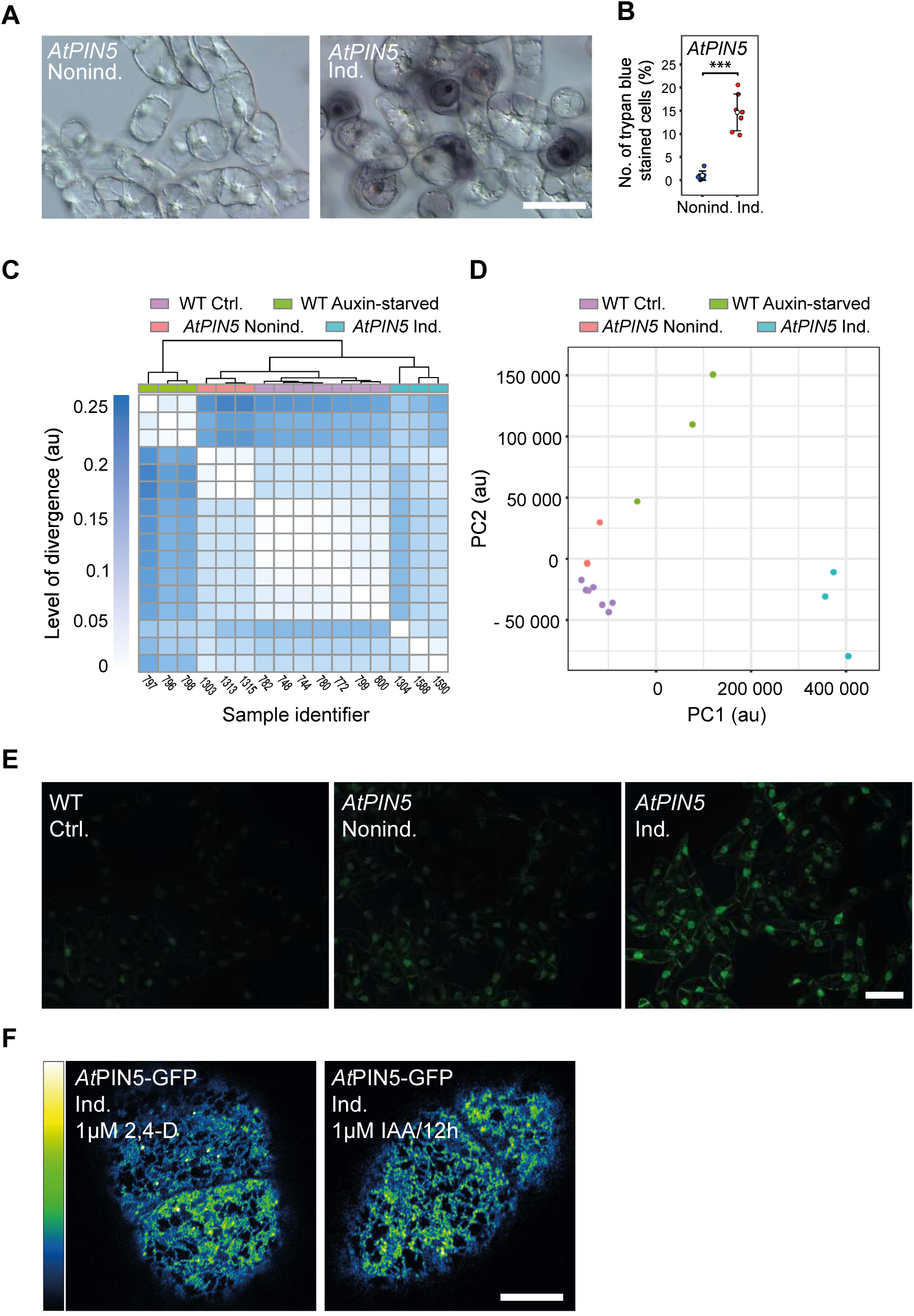
Increased frequency of symptoms of cell death and its relationship to auxin starvation following induction of *AtPIN5*. **A** and **B)** The representative images of incorporation of the trypan blue inside the cells following induction of *GVG::AtPIN5* **A)**, quantified on **B)**. **C** and **D)** Quality control of RNA-Seq data processing, a heat-map representing Jensen-Shannon divergence between individual samples (sorted by internal sample IDs) **C)**, projections of individual samples onto the principal components (PC) **D)**. **E)** An increased fluorescence on confocal images of the *GVG::AtPIN5* cells stained by 10 µM H2DCFDA to detect ROS, following the transgene induction. **F)** The pattern and intensity of the *At*PIN5-GFP foci are comparable in both standard 2,4-D-containing medium and that continuously supplemented with IAA every 12 hours. The signal intensity is indicated by the green fire blue rainbow lookup table on **F)**. On **B)**, the dot plot of a data set of 2000 cells in total, the open symbols are averages ± S.E., \*\*\**P* < 0.001 by Student’s *t*-test. Scale bars: 25 μm on **A)**, 100 µm on **E)**, and 10 µm on **F)**.

**Figure S3.**
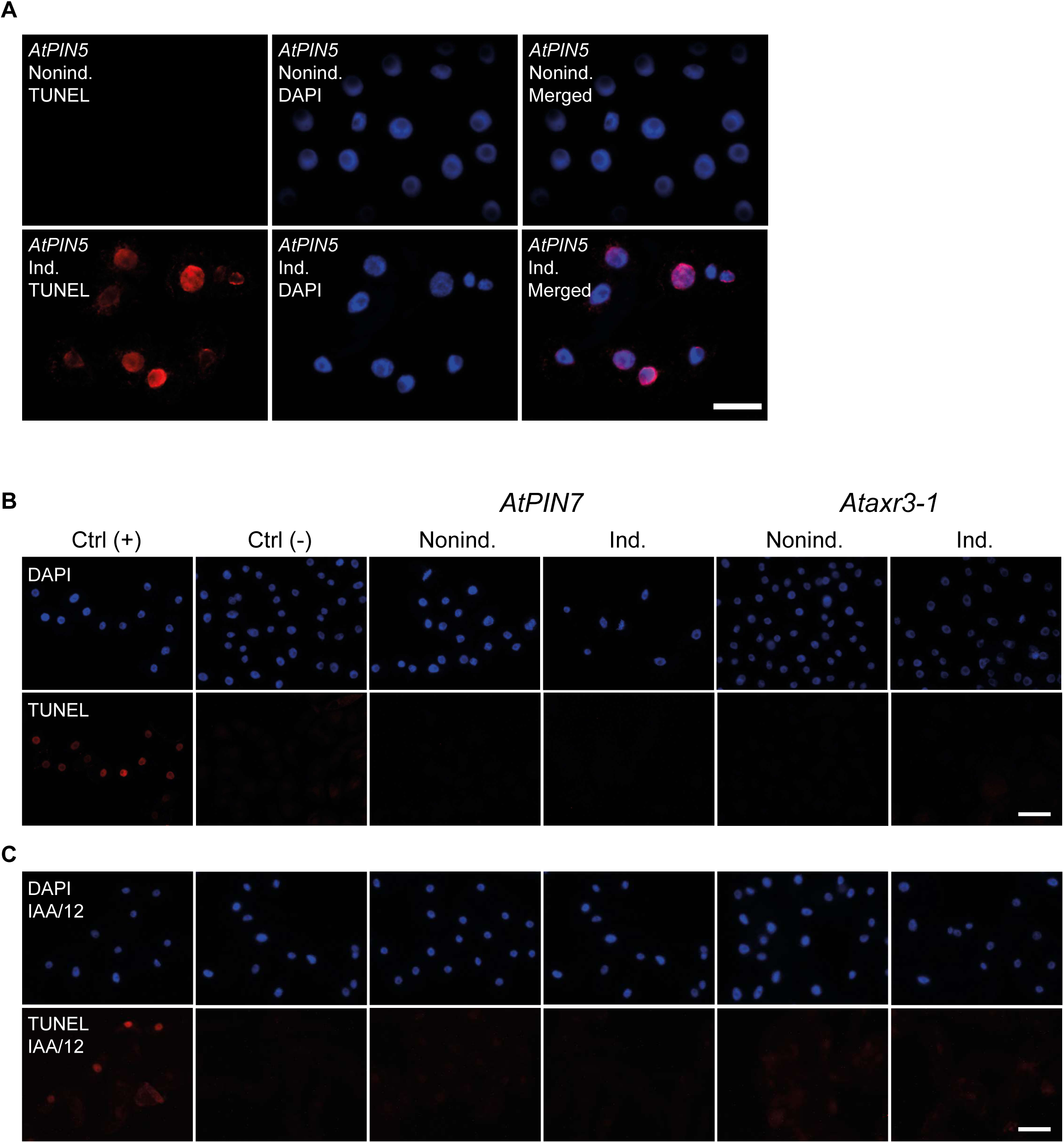
Detection of the DNA fragmentation during auxin deficiency. **A)** TUNEL staining (left: TUNEL, middle: control nuclear DAPI staining, right: merged false-colored images) marks an enhanced disintegration of nuclear DNA in the cells where *GVG::AtPIN5* was induced. **B** to **C)** Negative TUNEL stainings in *GVG::AtPIN7* and *XVE::Ataxr3-1* cells evidence the absence of the PCD, both under the standard conditions **B)**, or when supplemented with IAA every 12 hours during the monitored cultivation period **C)**. Upper panels: control nuclear DAPI staining, lower panels: TUNEL. Ctrl (+): control cells additionally treated with DNAse I to detect the positive TUNEL reaction, Ctrl (-): staining where the TdT enzyme action was omitted from the control reaction. Scale bars: 50 μm.

**Figure S4.**
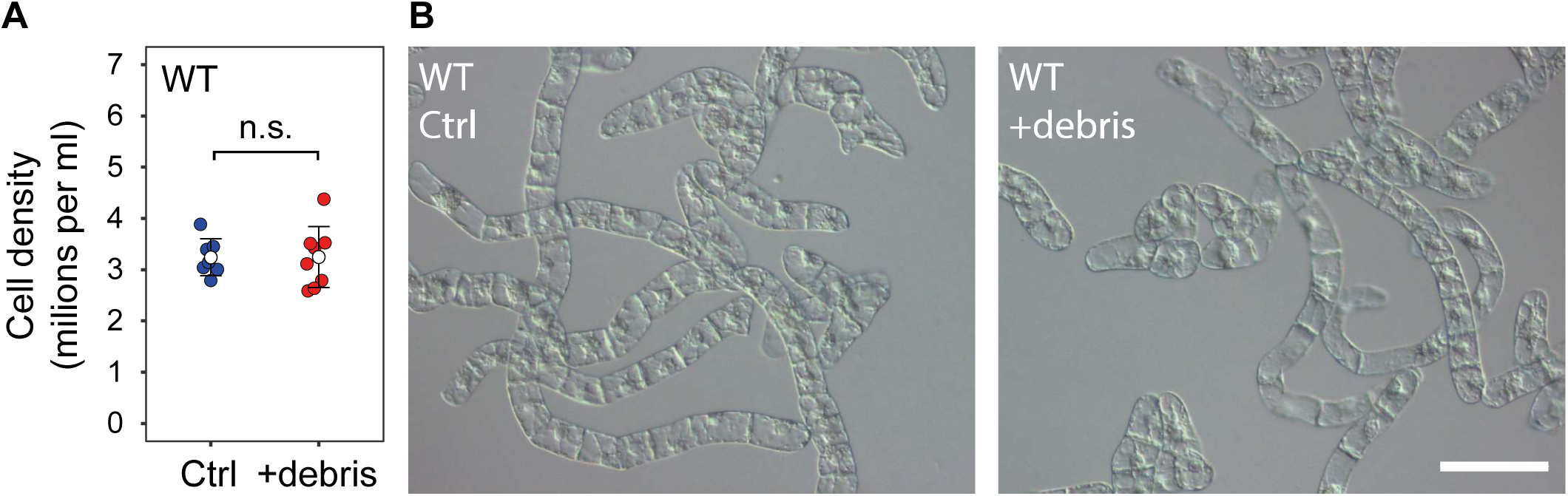
WT BY-2 cells cultured in a medium containing cell debris exhibit neither decrease of proliferation activity **A)** nor signs of decreased viability monitored by trypan blue staining **B)**. Scale bar: 100 μm on **B)**.

**Figure S5.**
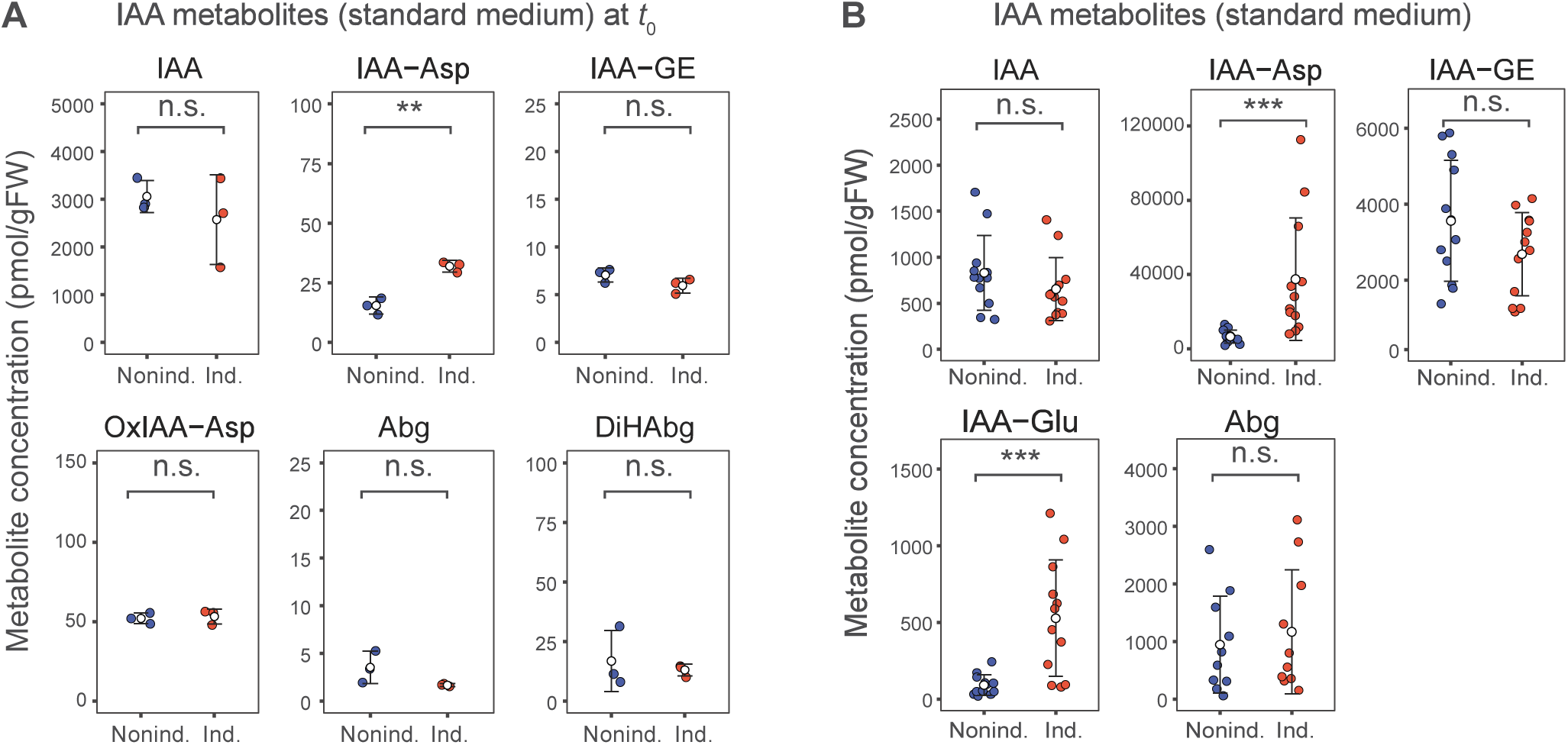
Additional experiments examining the activity of the IAA decarboxylation pathway following the *AtPIN5* expression. **A)** Representative LC-MS analysis of IAA and its metabolites in cells expressing *XVE::AtPIN5* determined immediately after 1 µM IAA supply (*t*_0_). While the exogenously applied IAA is up-taken immediately, the levels of its metabolites are still barely detectable. **B)** LC-MS analysis of IAA and its metabolites in cells expressing *GVG::AtPIN5* cultured on the standard 2,4-D-supplemented media, 2 hours after adding 1 µM IAA. Open symbols indicate the mean of three technical measurements; the bars are ± S.E. \*\**P* < 0.01, \*\*\**P* < 0.001 by Student’s *t*-test, n. s.: not significant at *α*_0.05_. IAA: indole-3-acetic acid, IAA-Asp: IAA-aspartate, IAA-GE: IAA-glucose ester, OxIAA-Asp: Oxo-IAA-aspartate, Abg: ascorbigen, DiHAbg: dihydroascorbigen.

**Table S1.** Results of PANTHER Overrepresentation test (GO - Molecular function) of transcripts significantly upregulated in both auxin-starved and the *XVE::AtPIN5*-expressing cells.

**Table S2.** Results of PANTHER Overrepresentation test (GO - Molecular function) of transcripts significantly upregulated in BY-2 cells carrying induced *XVE::AtPIN5*, compared to untreated controls and auxin-starved cells.

